## Supplementary Information for "AlphaFold-Metainference: Prediction of Structural Ensembles of Disordered Proteins"

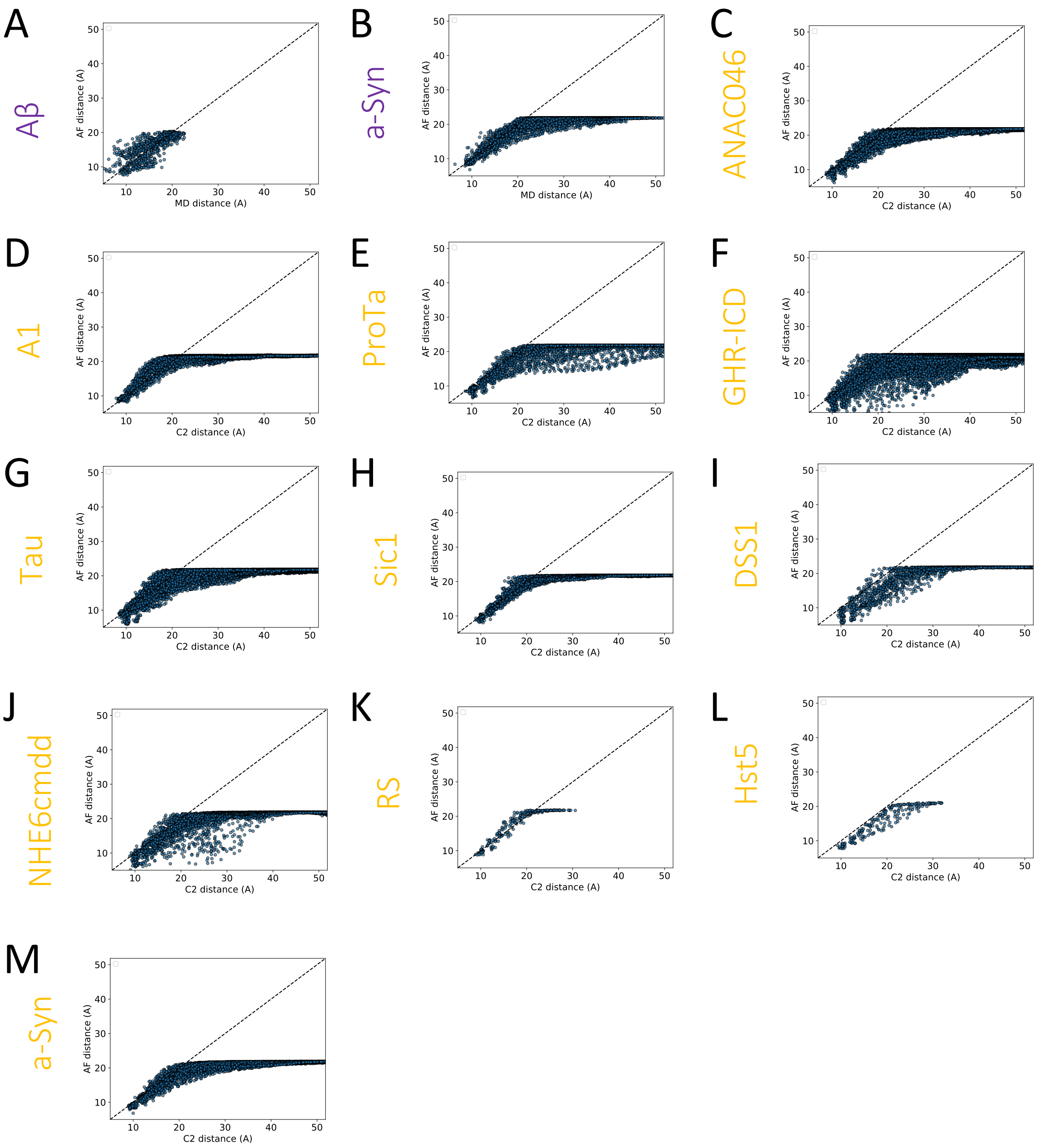


**Figure S1. Comparison of average inter-residue distances predicted by AlphaFold and back-calculated from molecular simulations. (A,B)** Distances from all-atom molecular dynamics (MD) simulations. **(C-M)** Distances from CALVADOS-2 (C2) simulations.

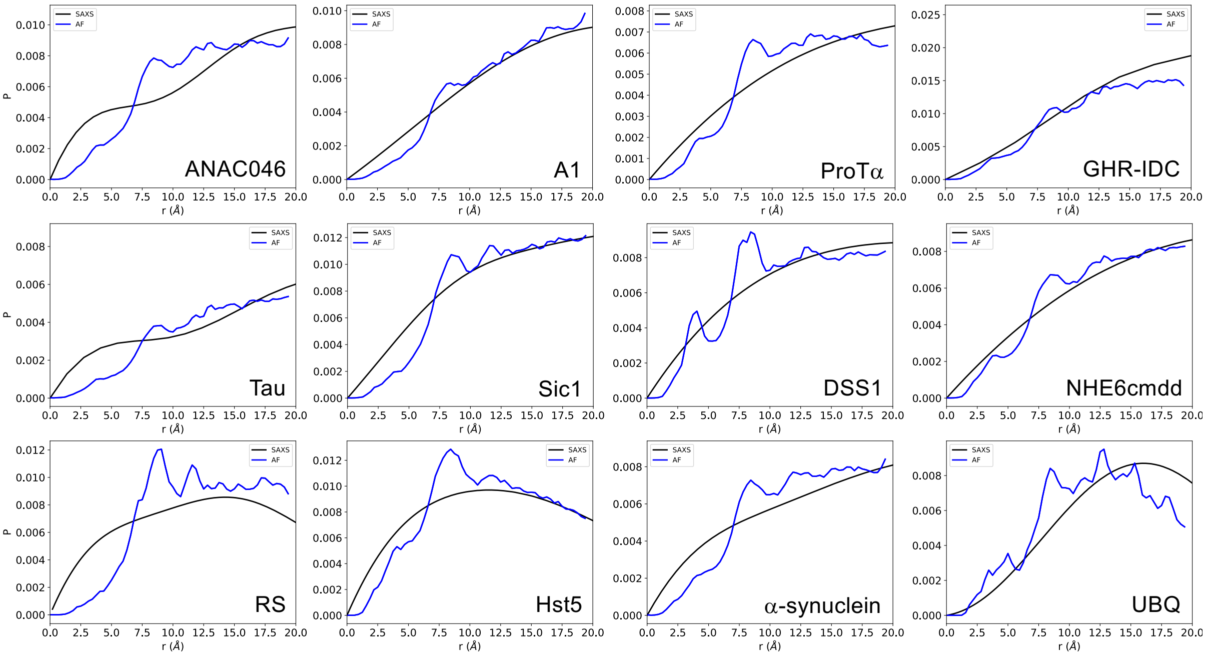


**Figure S2. Comparison of inter-residue distance distributions obtained by SAXS and predicted by AlphaFold.** We used a set of 11 proteins for which both SAXS and NMR diffusion measurements were available[1]. SAXS-derived inter-residue distance distributions are shown in black, and AlphaFold-predicted average inter-residue distance distributions are shown in blue. The cut-off distance in the AlphaFold predictions is 21.84 Å.

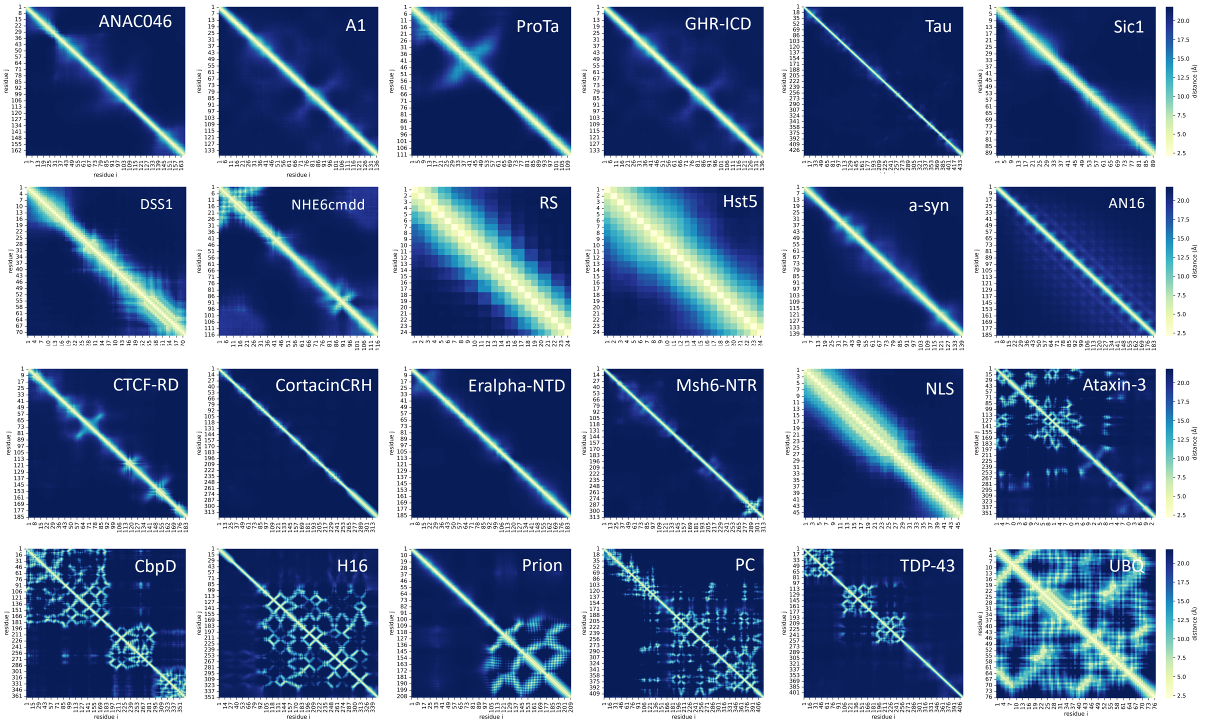


**Figure S3. AlphaFold distograms of the proteins analyzed in this work.** The distograms report the average inter-residue distances predicted by AlphaFold for the intrinsically disordered proteins, partially disordered proteins and folded proteins that we reported in this work.

A

| Protein | AF-MI | AF | Exp. Rg (ATSAS) [1] | Length | v |
|---|---|---|---|---|---|
| ANAC046 | 3.63 | 3.16 | 3.6 ± 0.3 | 167 | 0.55 |
| A1 | 2.65 | 2.72 | 2.5 ± 0.1 | 137 | 0.49 |
| ProTa | 4.14 | 3.03 | 3.7 ± 0.2 | 111 | 0.62 |
| GHR-ICD | 6.4 | 3.84 | 6.0 ± 0.5 | 351 | 0.56 |
| Tau | 7.18 | 4.71 | 6.4 ± 0.5 | 441 | 0.54 |
| Sic1 | 2.85 | 2.8 | 2.9 ± 0.1 | 90 | 0.58 |
| DSS1 | 2.6 | 2.06 | 2.5 ± 0.1 | 71 | 0.58 |
| NHE6cmdd | 3.12 | 2.74 | 3.2 ± 0.2 | 116 | 0.57 |
| RS | 1.34 | 1.72 | 1.26 ± 0.08 | 24 | 0.57 |
| Hst5 | 1.36 | 1.57 | 1.34 ± 0.05 | 24 | 0.58 |
| a-syn | 3.64 | 2.93 | 3.56 ± 0.04 | 140 | 0.56 |

B

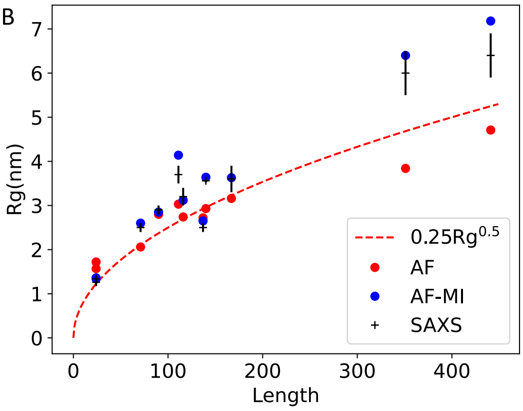


**Figure S4. Comparison of the Rg values from SAXS measurements with those from the AlphaFold (AF) and AlphaFold-Metainference (AF-MI) simulations. (A)** Rg values predicted by AlphaFold-Metainference, AlphaFold individual structures and SAXS, as well as length and scaling exponent ν. **(B)** Rg as a function of protein length. We report predictions by AlphaFold-Metainference (blue), with AlphaFold individual structures (red) and SAXS data (black)[1]. The theoretical scaling of Rg with the length for a Flory random coil (ν=0.5) is shown as a dotted red line.

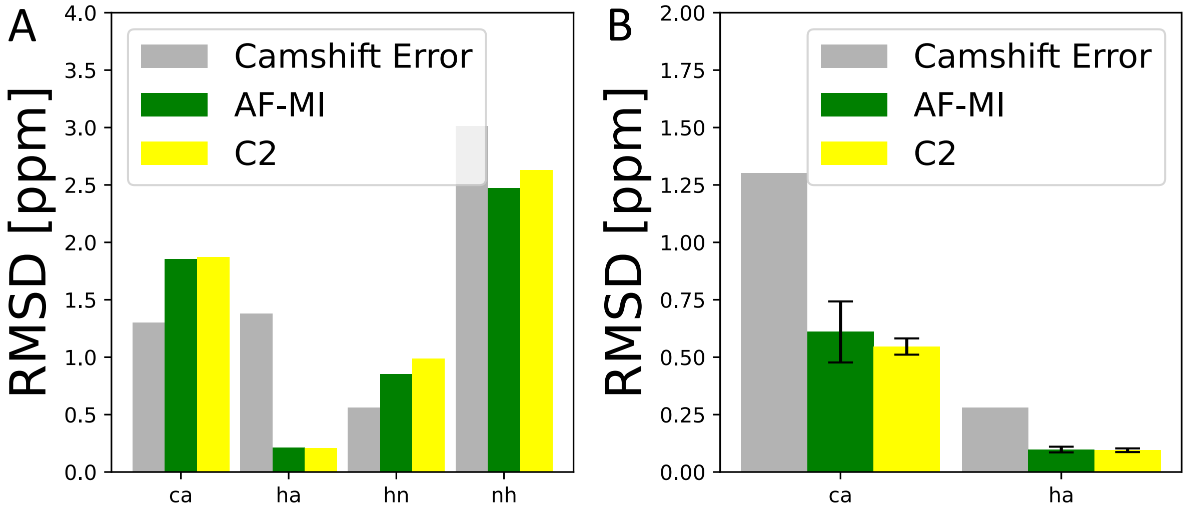


**Figure S5. Comparison of the back-calculated NMR chemical shifts from the CALVADOS-2 and AlphaFold-Metainference simulations. (A,B)** Root mean square deviation (RMSD) between CA, HA, HN, and NH chemicals shifts from Refs.[3,4] and predicted[5] ones from AlphaFold-Metainference (AF-MI, green) and CALVADOS-2 (C2) ensembles (yellow) for Sic1 (A) and AN16 (B). The errors of the CamShift chemical shift predictions are shown as grey bars.

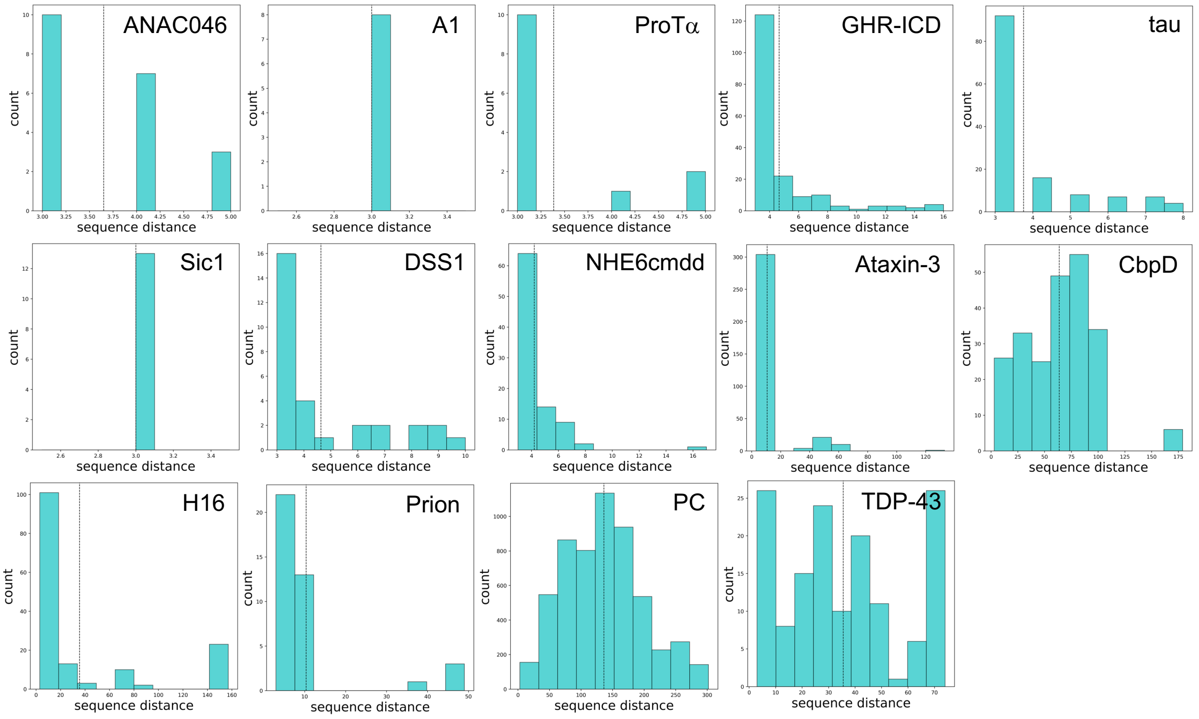


**Figure S6. Sequence separation of the pairwise distances used as structural restraints in the AlphaFold-Metainference simulations.** The histograms show that distance along the sequence (i.e. the sequence separation) of residue pairs used as structural restraints of the highly disordered proteins shown in **Figures 1-2** tend to be small, while those of the partially disordered proteins shown in **Figures 3-6** can be quite large. The dotted lines indicate the average sequence separation of pairwise distances used as restraints per system.

**A**

| | ΔRg/Rexp % | | | | | | ΔRg/Rexp % |
|---|---|---|---|---|---|---|---|
| System | PAE<5 | PAE<7 | PAE<10 | PAE<20 | Hydropathy | Plddt>75 for >5 residues | Calvados |
| AN16 | -15.60 | -15.93 | -18.74 | -14.52 | -0.98 | | -24 |
| NLS | -35.83 | -22.76 | -22.92 | -23.33 | -2.28 | | -21 |
| CTCF-RD | 0.22 | 21.76 | 17.58 | 13.44 | -0.73 | | 15 |
| Eralpha-NTD | 9.85 | 42.23 | 41.30 | 12.67 | -0.57 | | 18 |
| Msh6-NRT | -33.04 | -9.63 | -1.96 | 15.00 | -1.417 | | -22 |
| CortactinCRH | 18.78 | 51.20 | 31.01 | 52.74 | -1.2 | | 14 |

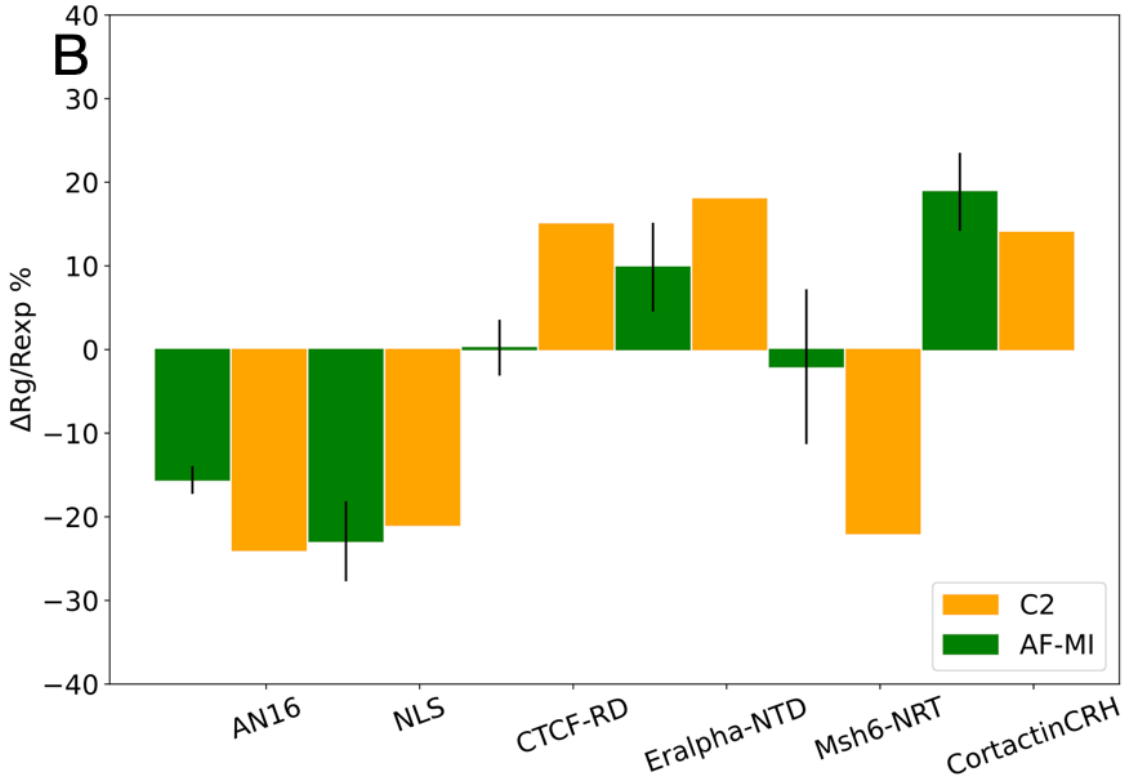


**Figure S7. Selection criterion for AlphaFold-predicted distances. (A)** Agreement with experimental Rg values of the AlphaFold-Metainference structural ensembles using different PAE values (<5, <7, <10, <20) for a set of benchmark proteins from Ref[2]. Proteins meeting the criterion of hydropathy < -1.4 and at least a 5-residue tract with pLDDT>75 are shown in green (PAE<10), while the remaining proteins are shown in red (PAE<5). **(B)** Comparison of the agreement between experimental Rg values and those back-calculated from the AlphaFold-Metainference structural ensembles, using the distance selection criterion reported above, as well as those back-calculated from the CALVADOS-2 structural ensembles.

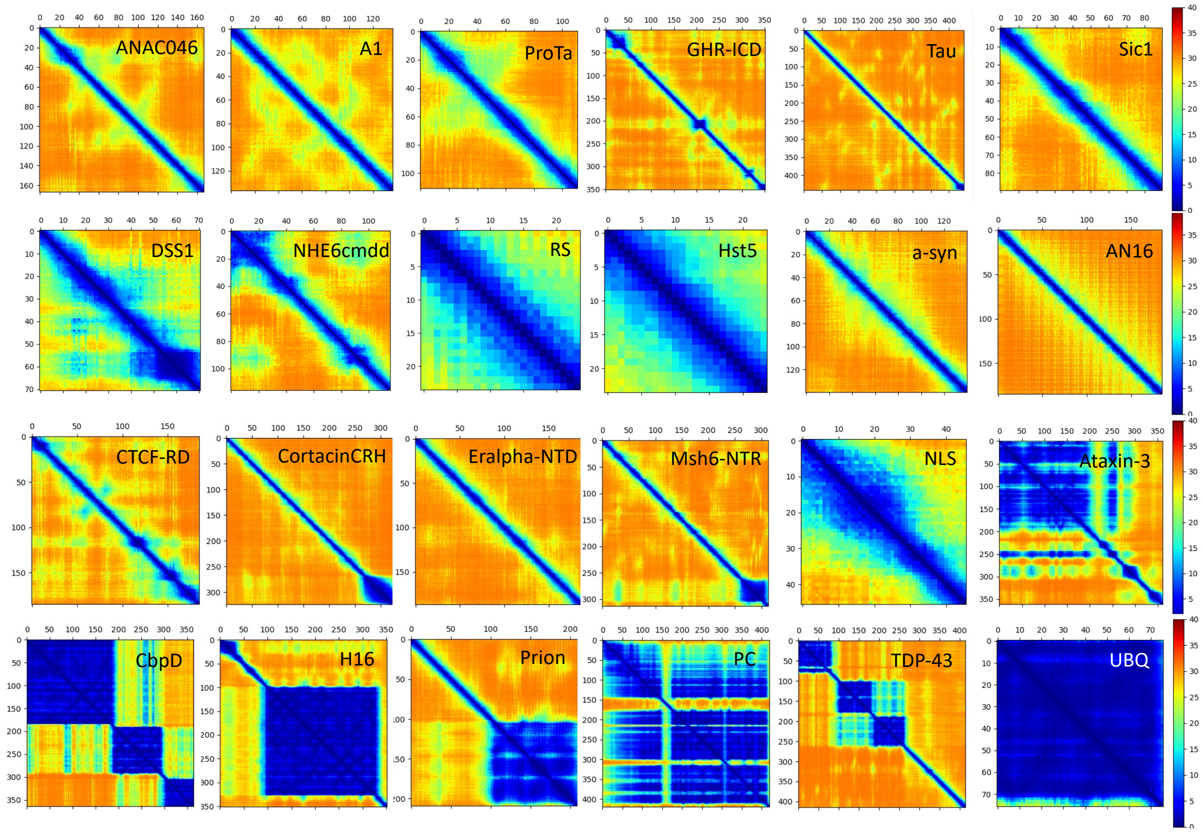


**Figure S8. AlphaFold PAE maps for the proteins analyzed in this work.** The maps report the predicted aligned error (PAE) of AlphaFold for the intrinsically disordered proteins, partially disordered proteins and folded proteins that we used in this work.
